# Vitamin B12 Improves Skeletal Muscle Mitochondrial Biology in Aged Mice

**DOI:** 10.64898/2026.01.13.699119

**Authors:** Abigail R. Williamson, Angad Yadav, Wenxia Ma, Susan Schmitt, Shelby Rorrer, Lauren E. Odom, Luisa F. Castillo, Chloe T. Purello, Olga V. Malysheva, James Mobley, Martha Field, Anna E. Thalacker-Mercer

## Abstract

Age-related skeletal muscle deterioration is a commonly reported disability among older adults, attributed to several factors including mitochondrial dysfunction, a major hallmark of aging. Therapies to attenuate or reverse mitochondrial decline are limited. Despite identified positive relationships between vitamin B12 (B12) and mitochondrial biology, the impact of B12 supplementation on skeletal muscle mitochondria, in advanced aged, has not been examined. Thus, the impact of B12 supplementation on skeletal muscle mitochondrial biology was examined in (i) aged female mice, given 12 weeks of B12 supplementation (SUPP) or vehicle control, and (ii) in human primary myotubes. In the mouse model, mitochondrial DNA and content were measured with PCR and citrate synthase activity, respectively; mitochondrial morphology was examined using transmission electron microscopy; mitochondrial function was examined using extracellular metabolic flux analysis; and proteins and pathway enrichment was identified with proteomics. In the cell model, ROS and glutathione was measured using luminescent assays. The results demonstrated that SUPP in aged mice increased muscle mitochondrial content and improved morphology. Further, differentially expressed proteins were enriched in TCA cycle, OXPHOS, and oxidative stress pathways. In the cell model, B12 supplementation reduced ROS levels. This is the first study, to our knowledge, examining the impact of B12 supplementation on skeletal muscle mitochondrial biology in aged female mice. Results suggest that B12 supplementation improves mitochondrial biology in aged female mice.

## INTRODUCTION

The age-related deterioration of skeletal muscle mass, function, and strength (i.e. sarcopenia) is a common phenotype of aging, associated with decreased quality of life, physical disability, and mortality (1, 2). Despite the growing population of adults over the age of 65 y, to date, treatment(s) to attenuate and/or reverse the characteristics of sarcopenia are minimal and consist primarily of exercise (3–5); thus, expansion of therapeutic options to reverse or attenuate muscle decline is necessary.

Studies have demonstrated that the micronutrient cobalamin (i.e., vitamin B12 [B12]) is associated with indicators of sarcopenia. For example, low B12 status, which can be due to reduced dietary intake and absorption as well as medication and alcohol use (6), has been associated with decreased skeletal muscle mass, strength, and quality in older adults (7–10). B12 supplementation, on the other hand, has been shown to improve muscle quality and strength in B12 deficient older adults (11). How B12 availability impacts skeletal muscle mass, strength and function in older adults remains unclear.

Skeletal muscle represents a mitochondrially dense tissue, susceptible to age-related mitochondrial changes. Reduced oxidative phosphorylation (OXPHOS), mitochondrial content and mitochondrial number (12) characterize mitochondrial dysfunction and have been observed with sarcopenia (2, 13–15). Importantly, reduced mitochondrial content and function as well as disrupted morphology have been observed in non-muscle tissues with B12 deficiency in young animals (16–18). For example, in young mice, B12 deficiency (dietary and genetic deficiency) impaired liver mitochondrial complex I and IV activity (17). In a similar study, increased uracil was observed in liver mtDNA of B12 deficient mice, which was associated with decreased mtDNA content and impaired mitochondrial function (16). In young sheep, B12 deficiency led to distorted mitochondrial morphology, inclusive of visually enlarged mitochondrial size and impaired cristae structure, compared to B12 sufficient sheep (18). These morphological changes are similar to what is seen in mitochondria of aged mouse and human skeletal muscle tissue (13, 15). Despite the identified relationships between B12 availability and mitochondrial biology, the impact of B12 supplementation on skeletal muscle mitochondria, in advanced aged, has not been examined. Thus, the purpose of this study was to evaluate the impact of B12 supplementation on skeletal muscle mitochondrial biology in aged mice. Based on the limited available research, we hypothesized that B12 supports mitochondrial biology (i.e., content, function, and structure) and that supplementing aged mice with B12 would impact mitochondria biology in older adults.

## METHODS

### Animal Model

This study was approved by the Institutional Animal Care and Use Committee (IACUC-22721) at The University of Alabama at Birmingham (UAB). Aged female C57BL/6N mice (20-22 months old) were obtained from Charles River Laboratories, USA, and acclimated for 4 weeks before experimental interventions began. Female mice were selected as the primary sex for our models as previous studies examining vitamin B12 effects on mitochondrial function were conducted exclusively in males (15, 16), necessitating evaluation of sex-specific responses. 20-22 months old mice were selected due to onset of established aging biomarkers including reduced grip strength, endurance, and muscle mass characteristic of a sarcopenic phenotype that occur at ∼18 months (19).

All mice were maintained on a standard AIN93G diet with water and food provided *ad libitum* throughout the study duration. Mice were randomly assigned to one of two groups: a vehicle control group (V-CNT) and a B12-supplemented group (SUPP). Both groups received weekly intramuscular injections alternating between the tibialis anterior (TA) and gastrocnemius muscles. The V-CNT group received bilateral 30 μL saline (0.9% NaCl), while the SUPP group received bilateral 0.65 mg cyanocobalamin (Sigma Aldrich 47869) dissolved in 30 μL saline (0.9% NaCl), providing a total dose of 1.30 mg vitamin B12 per mouse per week for 12 weeks.

### Body weight and food intake assessment

Body weight was measured weekly throughout the 12-week treatment period for both V-CNT and SUPP groups using a calibrated digital scale (**SFig. 1A**). There was no significant difference in weight between treatment groups over the course of the twelve-week treatment (**SFig. 1A**). Mice were weighed at the same time of day to minimize circadian variation. Food intake was monitored weekly by weighing food before placing in food hoppers of cages and again at the end of each measurement period (**SFig. 1B**). Daily food intake per mouse was calculated by dividing the total food consumed by the number of days and the number of mice per cage. There was no difference in food consumption between treatment groups over the course of twelve-week treatment.

**Fig. 1.**
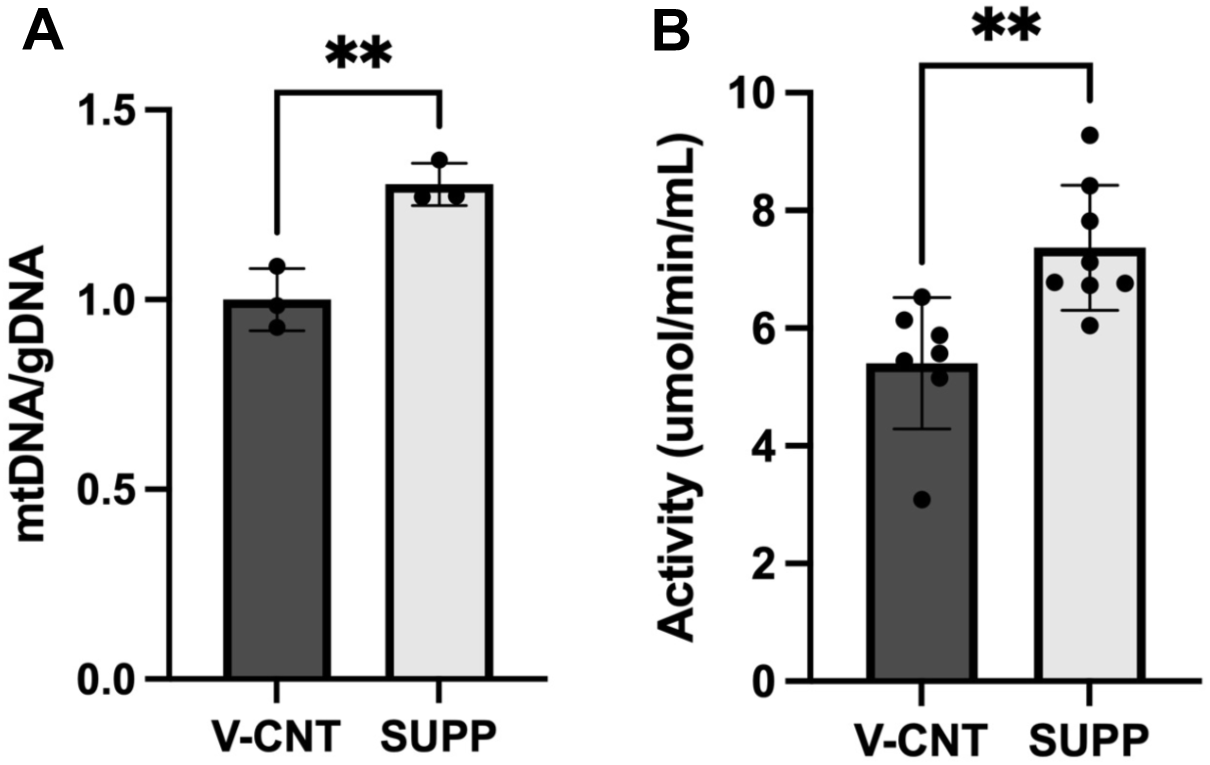
B12 supplementation improves mitochondrial DNA and content. **A.** mtDNA was greater in gastrocnemius tissue of B12 supplemented aged female mice (SUPP) compared to vehicle control treated mice (V-CNT) as measured by PCR and normalized to genomic DNA. **B.** Citrate synthase activity was greater in gastrocnemius tissue of SUPP compared to V-CNT. **p<0.01

### Tissue and blood collection and processing

Following the 12-week treatment period, TA quadriceps, gastrocnemius, and soleus muscles were harvested from each mouse (V-CNT n=8, SUPP n=10). Muscles were cleaned of excess connective tissue and fat, blotted dry, and processed for subsequent analyses. One TA was processed for transmission electron microscopy (TEM). Briefly, fresh TA was cut longitudinally into approximately 10 mg pieces and immediately fixed in 2% paraformaldehyde / 3% glutaraldehyde in 0.1 M sodium cacodylate buffer (pH 7.4). The remaining muscles were snap-frozen in liquid nitrogen and stored at -80°C for future analysis.

To measure treatment differences in muscle mass, at 12 weeks the muscle mass of the left quadricep was quantified and normalized with bone (femur) length. To do this, prior to freezing, after visible connective tissue and fat were removed from the muscle and the muscle blotted dry, the muscle was immediately weighed on a calibrated analytical balance and mass recorded. Femur length was measured in millimeter (mm) using a ruler. Muscle weights were normalized to bone length (g/mm) to account for variations in body size that would impact muscle mass. SUPP mice had increased muscle mass compared to V-CNT (**SFig. 1C**).

Blood was collected from each mouse in a non-heparin coated microcentrifuge tube at the time of sacrifice. Blood was then centrifuged at 1500xg for 15 minutes to obtain serum for serum methylmalonic acid (MMA) quantification. MMA is a biomarker of B12 status and was quantified using a Q Exactive Orbitrap mass spectrometer (Thermo Fisher Scientific), as previously described (19). All tissue and serum samples were stored at -80°C until needed.

### Real-Time qPCR Quantification

Mitochondrial DNA (mtDNA) content was determined using RT-qPCR. Approximately 30 mg of snap-frozen gastrocnemius muscle from V-CNT (n=6) and SUPP (n=6) mice was lysed, and total DNA was extracted using the Epoch GenCatch™ DNA Miniprep Kit (24–60050). mtDNA content was assessed using real time qPCR on the ThermoFisher QuantStudio3 system, as previously described (20). DNA concentration and purity were assessed by Nanodrop. Primers targeting mtDNA (mMito: forward 5’-CTAGAAACCCCGAAACCAAA-3’, reverse: 5’-CCAGCTATCACCAAGCTCGT-3’) and B-2-microglobulin, a marker for genomic DNA, as a housekeeper gene (mB2M: forward 5’-ATGGGAAGCCGAACATACTG-3’, reverse 5’-CAGTCTCAGTGGGGGTGAAT-3’) were used (20). mtDNA content was normalized to genomic DNA to determine relative mtDNA. Relative expression was calculated using the 2^^-ΔΔ^Ct method.

### Enzymatic assay

Snap frozen quadriceps muscle from V-CNT (n=7) and SUPP (n=7) mice were used for citrate synthase (CS) activity, a marker of mitochondrial content (21). Activity was measured spectrophotometrically using the Cayman MitoCheck® citrate synthase activity assay kit (#701040) according to the manufacturer’s protocol. Samples were assessed in duplicate, and activity values were normalized to protein concentration which was determined using Pierce BCA Protein Assay.

### Transmission Electron Microscopy

Fixed TA muscles from V-CNT and SUPP groups (n=3 per group) were embedded in fresh Embed 812 at the UAB High Resolution Imaging Facility. Sections were cut using the Leica EM UC7 ultramicrotome, collected on 200 mesh copper grids, and stained with 1% alcoholic uranyl acetate and Reynolds lead citrate. Sections were examined using a JEOL JEM-1400 electron microscope. Images were captured with AMT digital software and ultrastructural images were acquired at 2000x and 5000x magnification. Captured images were blinded and measured for cristae score, by two independent lab personnel. 200 mitochondria were scored across three mice from each treatment group. Both intermyofibrillar (IMF) and subsarcolemmal (SS) regions were imaged. Cristae morphology from the IMF region was scored on a 0-4 scale, where: 0 = no cristae with sharp definition; 1 = >25% but <50% have no cristae; 2 = >50% but <100% of the mitochondrion area have no cristae; 3 = many cristae present but irregular in shape; or 4 = regular cristae (22, 23). Personnel were provided an example of each of the scores, based on these criteria.

### Global untargeted proteomics

For proteomics, snap frozen gastrocnemius muscle samples, randomly selected from V-CNT (n=3) and SUPP (n=3) mice, were lyophilized and prepared for proteomics analysis. Global untargeted proteomics was conducted at the UAB Mass Spectrometry and Proteomics Shared Resource, as previously described (24). All protein extracts in these studies were attained using T-PER™ Tissue Protein Extraction Reagent (Thermo Fisher Scientific) and quantified using Pierce BCA Protein Assay Kit (Thermo Fisher Scientific). The list of peptide IDs, generated based on SEQUEST search results, were filtered using Scaffold (Proteome Software, Portland, OR). Scaffold filters and groups all peptides to generate and retain only high confidence IDs while also generating normalized spectral counts (N-SC’s) across all samples for the purpose of relative quantification. The filter cut-off values were set with a minimum peptide length of >5 AA’s, with no MH+1 charge states, with peptide probabilities of >80% C.I., and with the number of peptides per protein ≥2. Scaffold incorporates the two most common methods for statistical validation of large proteome datasets, the false discovery rate (FDR) and protein probability (25–27). The protein probabilities were set to a >99.0% C.I., and an FDR<1.0. Relative quantification across experiments were then performed via spectral counting (28, 29) and when relevant, spectral count abundances were then normalized between samples (30). Downstream analyses, including functional annotation and pathway discovery, were conducted using Qiagen Ingenuity Pathway Analysis (IPA) (QIAGEN, Redwood City, CA). Volcano plots and heat maps from proteomics data were generated using R Studio.

### Extracellular metabolic flux analysis

Briefly, mitochondria-enriched homogenates from frozen gastrocnemius tissue were isolated in mitochondrial assay buffer (MAS) buffer. Mitochondrial protein concentrations were determined using the Pierce BCA Protein Assay. Complex I and Complex IV maximal activity was assessed using the Seahorse XF4 assay, as previously described (31).

### Cell model

Female Human Skeletal Muscle Myoblasts (HSMM; Lonza) were revived and proliferated in Skeletal Muscle Cell Growth Medium (Ready-to-use; Lonza), consisting of Basal Medium (Lonza) and Supplement Mix (Lonza), according to the manufacturer’s instructions. Upon reaching approximately 80-85% confluence, cells were switched to Skeletal Muscle Cell Differentiation Medium (Ready-to-use; PromoCell), composed of Basal Medium (PromoCell), Horse serum (2%, Cat no., manufacturer) and Supplement Mix (PromoCell), to induce myotube formation. After 5 days of differentiation, myotubes were left untreated (Control) or treated with 0.9% saline (Vehicle) or with 20 µM cyanocobalamin dissolved in 0.9% saline (B12) for 48 hours before processing for redox assays.

### Redox assays

The cell model was implemented to assess reactive oxygen species (ROS) presence and glutathione cycling, providing insights into B12’s impact on mitochondrial oxidative stress under conditions where tissue snap-freezing of the mouse muscle may impair accurate assessment of real-time cellular redox dynamics (32).

To measure ROS, HSMM myotubes were cultured and differentiated in 96-well white walled plates and divided into three groups (n=6 per group): negative control (untreated, Control), vehicle control (0.9% saline, Vehicle), and B12-supplemented (20 µM cyanocobalamin diluted in 0.9% saline, B12). After 48 hours of treatment, cells were assessed for ROS using the ROS-Glo kit (G8820, Promega, USA) following the manufacturer protocol. Briefly, cells were washed with 1X PBS (Gibco) and H_2_O_2_ substrate (ROS-Glo kit) was added to each culture and incubated for 6 hours at 37°C and 5% CO2. Following this, the ROS-Glo detection solution was added and incubated for 20 minutes at room temperature, after which, the relative luminescence was measured.

To measure total glutathione, oxidized glutathione (GSSG), and the reduced glutathione (GSH) to GSSG ratio (GSH:GSSG), HSMM myotubes were cultured and differentiated in 24-well plates and divided into three groups (n=6 per group): negative control (untreated, Control), vehicle control (0.9% saline, Vehicle), and B12-supplemented (20 µM cyanocobalamin dissolved in 0.9% saline, B12). After 48 hours of treatment, total glutathione and GSSG were measured spectrophotometrically at 412 nm using the Promega GSH:GSSG-Glo kit (V6611), according to the manufacturer’s protocol. The GSH:GSSG ratio was calculated using the formula: [(net total glutathione RLU minus net GSSG RLU)/(net GSSG RLU)] x 2, where the factor of 2 accounts for the stoichiometry of 2 moles of glutathione produced per mole of GSSG and RLU is relative light unit. The GHS:GSSG is reported as an indicator of redox state.

### Statistical analyses

For animal studies, comparisons between V-CNT and SUPP groups were performed using unpaired Student’s t-tests for normally distributed data, including average body weight, average food intake, muscle mass, TFAM protein level, citrate synthase activity, mtDNA content, Seahorse assay parameters. Normality was assessed using the Shapiro-Wilk test. Body weight and food intake, measured weekly over the 12-week treatment period, were analyzed using two-way repeated measures ANOVA with treatment (V-CNT vs. SUPP) and time (weeks) as independent factors, followed by Sidak’s post hoc test for multiple comparisons.

Mitochondrial cristae morphology was scored on a 0-4 ordinal scale, with higher score indicating better structural integrity (22, 23). To determine whether treatment altered the overall distribution of cristae quality, a t test was conducted for average cristae score per treatment group. To assess differences in cristae score within treatment groups, a one-way ANOVA was conducted with multiple comparisons.

For cell culture experiments (ROS and glutathione assays), one-way analysis of variance (ANOVA) followed by Tukey’s post hoc test was used to compare differences among three groups (Control, Vehicle, and B12).

All data are presented as mean ± standard deviation (SD). Statistical analyses were performed using GraphPad Prism (version 10)/R Studio (2022.12.0+353). Statistical significance was set at p<0.05 for all analyses. Sample sizes for each experiment are indicated in the respective methods sections and figure legends.

## RESULTS

### Intramuscular B12 injections improves B12 status in aged mice

There was a trend toward lower serum MMA (p=0.056), a functional marker of B12 status, in SUPP mice (vs. V-CNT mice), indicating greater B12 availability with B12 supplementation (**SFig. 2**).

**Fig. 2.**
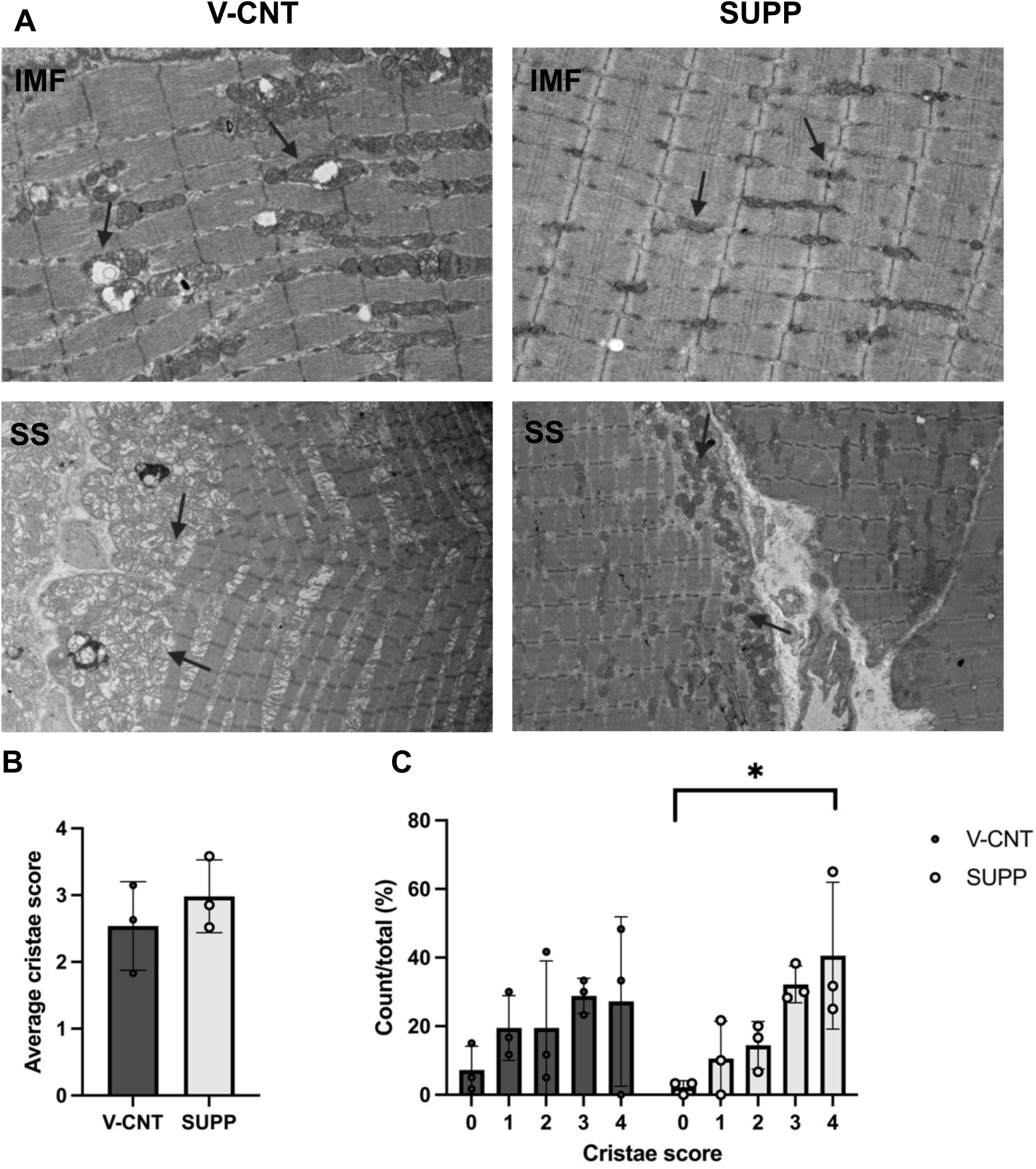
Higher prevalence of regular cristae in skeletal muscle mitochondria of B12 supplemented mice. **A.** Representative transmission electron microscopy images from female aged tibialis anterior muscle tissue, imaged for mitochondrial morphological assessment (n=3 per treatment). Vehicle treated mice (V-CNT) is displayed on the left, B12 supplemented (SUPP) on the right. IMF: intermyofibrillar, SS: subsarcolemmal. Visual morphological improvement can be seen among the supplemented tissue, including better cristae structure. Arrows indicate mitochondria. IMF images were captured at 5000x magnification. SS images were captures at 2000x magnification. **B.** Average overall cristae score within the IMF region (n=200 mitochondria, in each of n=3 mice) between treatment groups. **C.** Cristae score distribution was measured in IMF region in control (V-CNT) and treated (SUPP) groups. Data are shown as percentage of mitochondria per score. Statistical comparison was performed using a one-way ANOVA with multiple comparison test. For B and C the cristae score for each mitochondria measured was determined with the following criteria: 0 = no cristae with sharp definition; 1 = >25% but <50% have no cristae; 2 = >50% but <100% of the mitochondrion area have no cristae; 3 = many cristae present but irregular in shape; or 4 = regular cristae (23). *p<0.05

### B12 increases mitochondrial DNA and content

MtDNA and mitochondrial content are closely related to and used as indicators of mitochondrial function (21, 33). Increased mitochondrial content could further indicate an increase in mitochondrial biogenesis, which is typically seen to decline with advancing age, contributing to the loss of function in skeletal muscle (34). We assessed if mtDNA and CS were impacted by B12 supplementation. CS activity is a common marker for mitochondrial content (21). We observed increased (p=0.006) mtDNA in SUPP mice (vs. V-CNT mice) (**Fig. 1A**); mtDNA has previously been observed to decline with B12 deficiency (17). Additionally, CS activity was significantly higher (p=0.0048) in the muscle of SUPP mice compared to V-CNT mice (**Fig. 1B**).

### B12 improves mitochondrial morphology

As discussed, a prior study reported impaired mitochondrial morphology in B12 deficient sheep (18). Therefore, we next looked for morphological differences of the mitochondria using TEM. In the muscle of SUPP mice, after 12 weeks of supplementation, we observed visually smaller, more compact, mitochondria within the IMF region compared to the enlarged mitochondria with disrupted cristae in the V-CNT mice (**Fig. 2A**). Visual morphological differences were also seen between SUPP and V-CNT mitochondria within the SS region. Larger mitochondria are typically associated with disease states, inclusive of cellular stress, and often result in further alterations to mitochondrial shape and dynamics (35).

To assess the impact of B12 on mitochondrial ultrastructure, cristae morphology was scored on a scale (0-4) for 200 IMF mitochondria in 3 different mice for each treatment group, based on defined characteristics. Mitochondria in SUPP muscle exhibited a clear shift toward higher cristae integrity compared with V-CNT, with a greater proportion of mitochondria displaying well-defined, densely packed cristae (score 4) and fewer mitochondria lacking or showing poorly formed cristae (score 0). In contrast, V-CNT tissue showed a distribution skewed toward lower and intermediate scores, consistent with reduced cristae organization. No significant difference was observed between treatment groups when the cristae scores for 200 mitochondria were averaged (**Fig. 2B**). However, assessing cristae score within treatment groups revealed a significant difference between percent mitochondria scoring a 0 and a 4 in the SUPP tissue, which was not seen within the V-CNT (**Fig. 2C**); SUPP tissue has significantly more mitochondria with a score of 4 compared to those that scored a 0, whereas V-CNT had no significant difference amongst any score categories (**Fig. 2C**). Collectively, these findings demonstrate that supplementation improves mitochondrial ultrastructural integrity, increasing the prevalence of highly organized cristae while reducing the proportion of structurally compromised mitochondria. Our findings align with mitochondria morphological observations in B12 deficient sheep (18) and suggest that B12 supplementation (i.e., greater B12 status) improves mitochondria morphology in aged female mice. Mitochondrial cristae are the site for mitochondrial oxidation and chemiosmosis, in which cristae organization is a direct regulator of mitochondrial health and efficiency (35).

### B12 supplementation impacts mitochondrial TCA and OXPHOS pathways

Because previous studies evaluating the effect of B12 status on mitochondria biology only evaluated the impact of B12 on mitochondria DNA, content, function and morphology, we wanted to assess the biochemical and/or molecular pathways impacted by B12. Therefore, we conducted proteomics on the skeletal muscle after 12 weeks of treatment. We demonstrated that 12 weeks of B12 supplementation altered the protein profile and molecular pathways in the gastrocnemius muscle, of aged female mice. There were 212 differentially expressed proteins (139 up and 73 down, **Fig. 3A**) between the SUPP and V-CNT female mice. Supporting changes we observed with mitochondrial DNA and content (CS activity), CS protein levels were increased 1.6-fold increased with B12 supplementation. Mitochondrial transcription factor A (TFAM), which is crucial for the maintenance of mtDNA copy number (36), mitochondrial biogenesis, and OXPHOS regulation (37) was also significantly enriched in SUPP (2.2-fold, p=0.02). Interestingly, TFAM can protect against ROS-induced mtDNA degradation (37), further supporting improved mtDNA health.

**Fig. 3.**
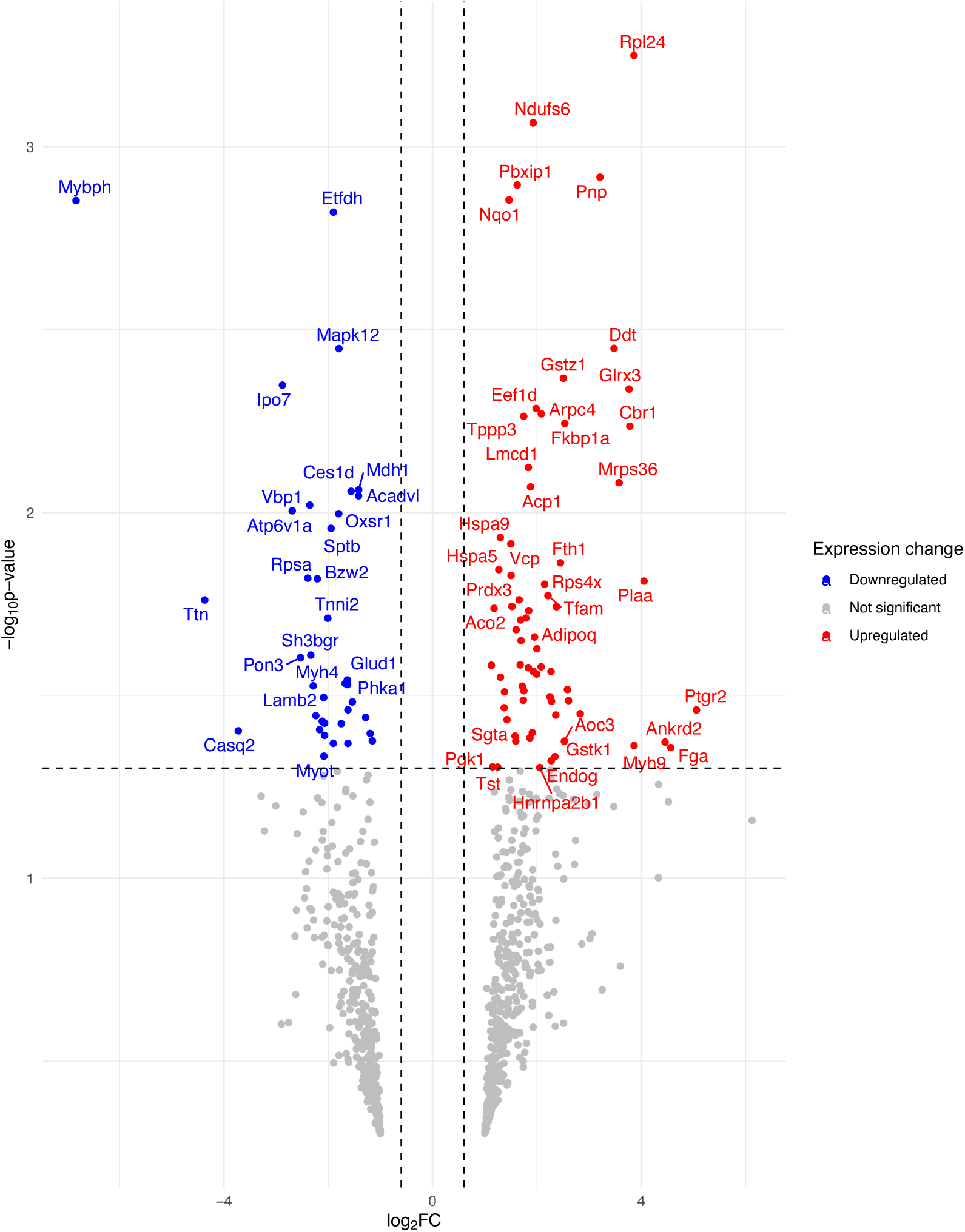
B12 Supplementation modifies the skeletal muscle proteome in aging. Volcano plot depicting differentially expressed proteins (n=211 proteins) in supplemented (SUPP) vs vehicle treated (V-CNT) gastrocnemius muscle; log_2_ fold change (x-axis) vs. -log_10_(p-value) (y-axis). Upregulated proteins are shown in red (139 total), downregulated in blue (73 total).

Next, we used IPA for functional annotation and network discovery of differentially expressed proteins. Further demonstrating that B12 availability impacts skeletal muscle mitochondria, we identified 43 mitochondria-related proteins that were differentially expressed between the SUPP and V-CNT female mice after 12 weeks of treatment (**Fig. 4A**). Intriguingly, one of the top canonical pathways, enriched with differentially expressed proteins, was *mitochondrial dysfunction,* which was downregulated in SUPP (z=-1.6, p=8.36E-11) (**Fig. 4B**). Additionally, in SUPP mice, the *respiratory electron transport (*z=0.8, p=1.21E-05), *TCA cycle* (z=1.0, p=2.09E-05), and *OXPHOS* (z=1.0, p=1.78E-03) canonical pathways were enriched with differentially expressed mitochondria proteins. These pathways included increased levels of isocitrate dehydrogenase [nad] subunit gamma 1, mitochondrial (IDH3G [1.8-fold]), CS (1.6-fold) and L-2-hydroxyglutarate dehydrogenase (L2HGDH [1.7-fold]). Additionally, proteins related to the electron transport chain, including complex I (NDUFA12 [2.7-fold], NDUFS6 [1.9-fold], NDUFS7 [4.3-fold]) and complex IV (COX5A [1.7-fold]) were enriched in SUPP. These findings suggest that B12 availability may impact complex I and IV activity (11, 38); therefore, we next tested whether B12 supplementation impacts mitochondria complex activity in aged female mice. Despite prior observations in the aged male mice, following 8 weeks of B12 supplementation (19), we did not observe an improvement in maximal oxygen consumption rate (OCR) at either complex I or IV with 12-weeks of B12 supplementation in aged female gastrocnemius (**SFig. 3**).

**Fig. 4.**
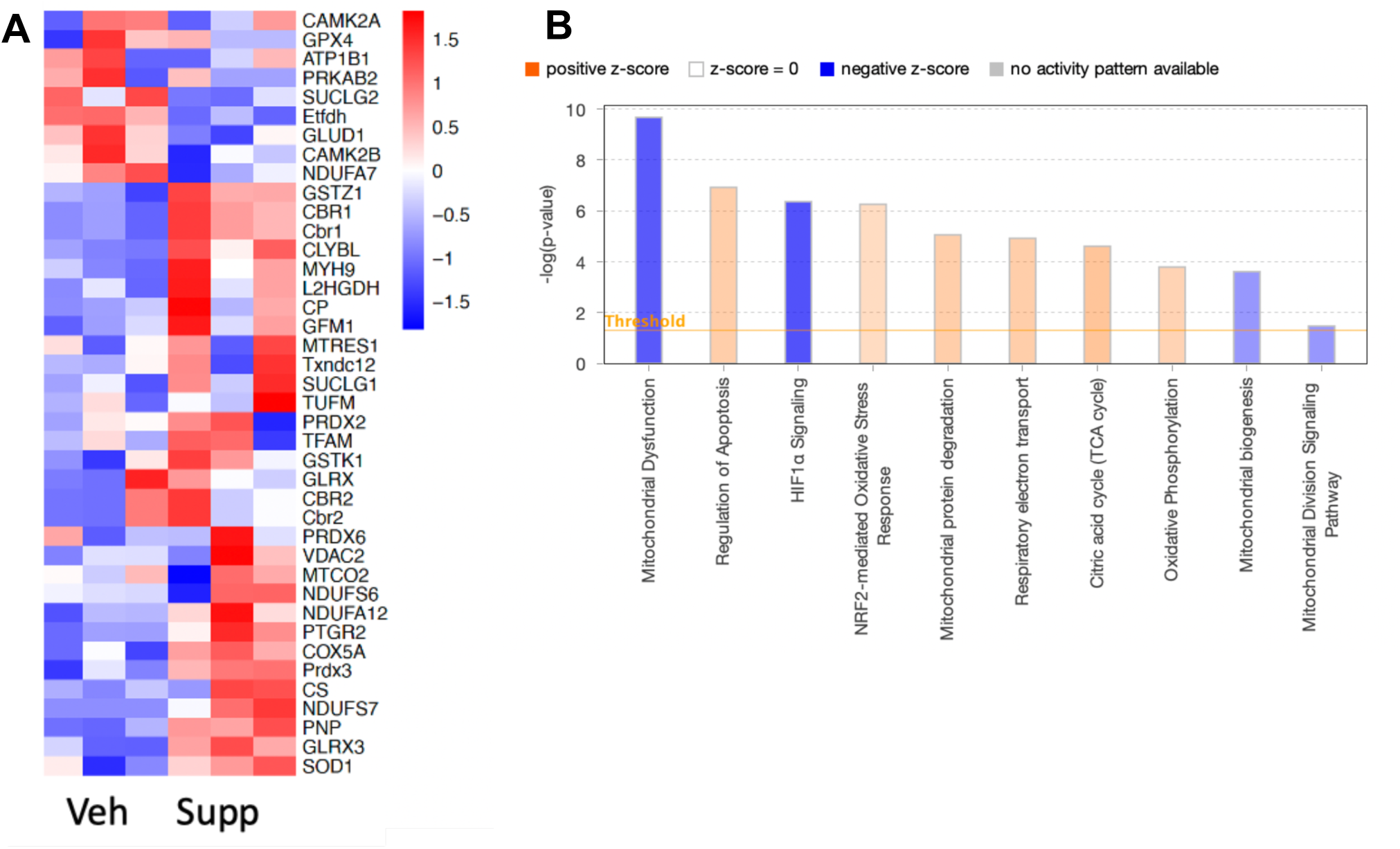
B12 supplementation leads to enrichment of mitochondrial proteins and pathways. **A.** Heat map displays mitochondria-related proteins based on fold change and adjusted p-value (p-value<1.0) in supplemented (SUPP) vs vehicle treated (V-CNT) mice. **B.** Top enriched pathways based on amount of related differentially expressed proteins presented in A. An enrichment of proteins that are classically related to mitochondrial dysfunction were observed within SUPP tissue, importantly, reflecting a downregulated pathway

### B12 impacts mitochondrial oxidative stress

Along with proteins related to mitochondrial metabolism, we also observed an enrichment of proteins related to oxidative stress regulation and protective mechanisms against oxidative damage (e.g., *NRF-2 mediated oxidative stress response*, z=0.80, p=5.58E-07, **Fig. 4B**) in SUPP tissue. In non-muscle cells, methylcobalamin, the active form of B12, has been shown to mitigate oxidative stress and protect against H_2_O_2_ oxidative stress through activation of the Nrf-2 pathway (39). Likewise, IPA downstream analysis predicted a decrease in the synthesis of ROS in SUPP mice (z=-1.745). Complimentary to this, one of the top-upregulated canonical pathways was *glutathione redox reactions* (p=0.013). Glutathione proteins play a role in detoxification of ROS and protect against oxidative damage (40). Glutathione S-transferase kappa 1 (GSTK1), a mitochondrial oxidative stress protein (41), was >2.0-fold higher in SUPP mice compared to V-CNT. Decreased GTSK1 levels are associated with increased oxidative stress (40).

Additionally, the copper transporter, ceruloplasmin (CP), was 4.3-fold increased with B12 supplementation, the largest fold-change among the differentially expressed proteins. Copper is a known antioxidant and shows antioxidant capability in skeletal muscle membrane (42). Intriguingly, CP has been seen to mimic B12 availability, suggesting it is a potential indicator of B12 status (43). Along with CP, the proteomics data demonstrated increased expression of several non-clinical indicators of improved B12 status including complement C3. Low C3 has been associated with B12 deficiency and that treatment with B12 supplementation improves C3 levels (44).

Recognizing observed changes in oxidative stress and antioxidant proteins, including glutathione proteins, we next questioned the impact of B12 supplementation on ROS and glutathione cycling. To address these questions, we conducted ROS and glutathione redox assays on human primary myotubes (i.e., differentiated HSMM cells) treated with and without B12 for 48 h. We demonstrated that, indeed, B12 supplementation (vs. Vehicle and Control [media alone]) reduced ROS levels in human myotubes (**Fig. 5A**). Additionally, in the glutathione assay, we did not observe a change in GSH:GSSG ratio with B12 supplementation in the primary human myotubes (**Fig. 5B**). Our primary cell culture results support the *in vivo* skeletal muscle proteomics and suggest that B12 supplementation reduces muscle oxidative stress via decreasing ROS levels, possibly through impacting ROS production and/or scavenging through routes other than glutathione. A reduction in ROS production in the muscle could underly observed improvements in mitochondrial morphology in the SUPP mice.

**Fig. 5.**
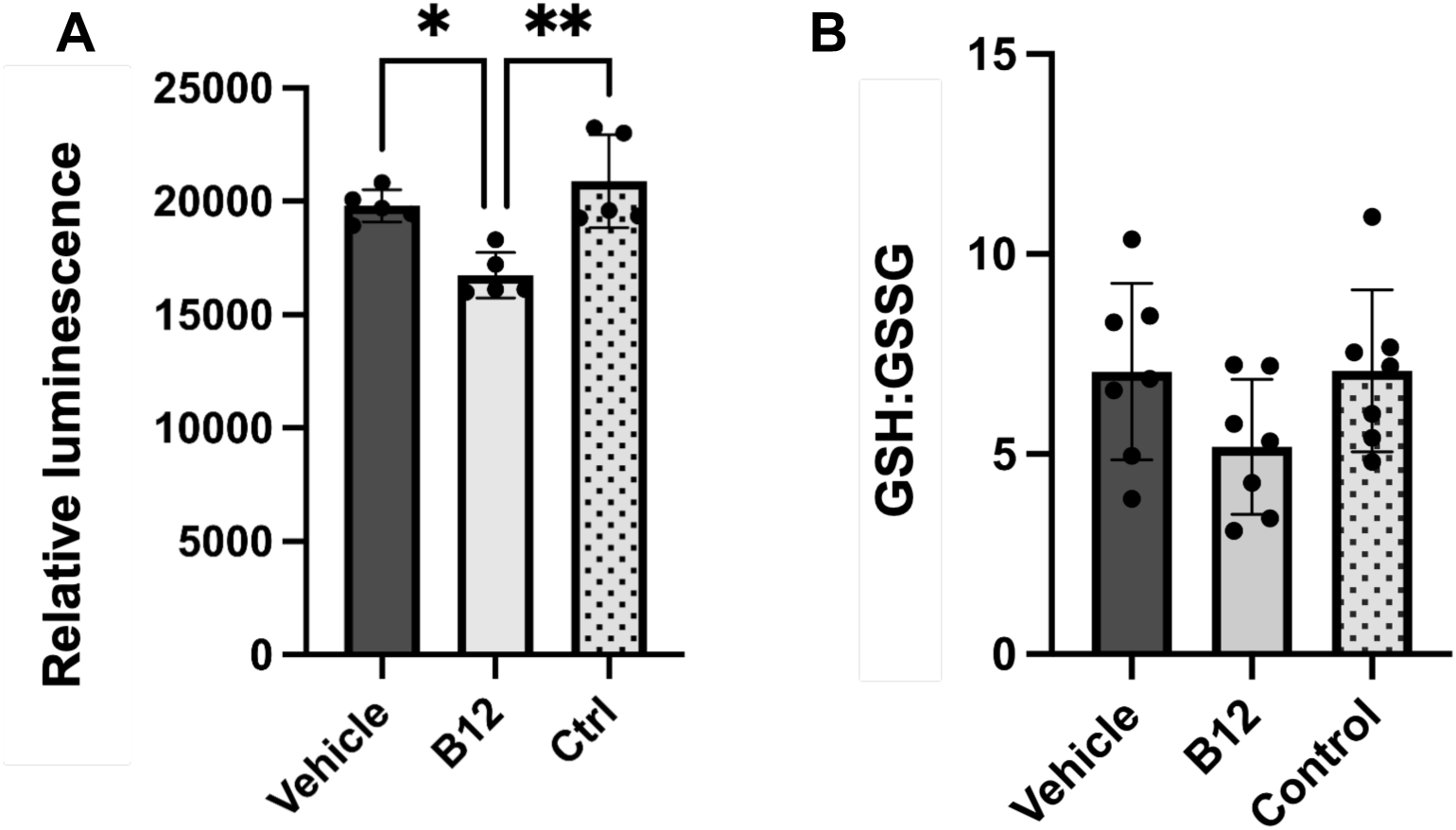
B12 supplementation reduces ROS production without altering glutathione levels in primary myotubes. **A.** ROS levels detected in female primary myotubes (n=6/group). ROS levels were significantly decreased in B12 supplemented (B12) female primary myotubes compared to vehicle treated (Vehicle) and control myotubes (Control). Cells were treated for 48 hours. **B.** Glutathione levels were unaffected by B12 supplementation in female primary myotubes compared to both Vehicle and Control. *p<0.05, **p<0.01

## DISCUSSION

This was the first study to demonstrate that B12 supplementation impacts skeletal muscle mitochondrial biology in aged female mice. B12 is water soluble, nutritionally essential micronutrient required for two intracellular enzymes (methylmalonyl-CoA mutase [MMUT] and methionine synthase [MTR]), whose metabolic pathways are relevant to mitochondria metabolism (directly and indirectly, respectively). However, the impact of B12 supplementation on mitochondria biology and the mechanism(s) in which B12 affects skeletal muscle have not been elucidated. This knowledge is important due to the relationship between B12 availability and characteristics of sarcopenia (7, 8). Our study demonstrated that B12 supplementation improved skeletal muscle mitochondria content and morphology in aged female mice. Further, our proteomics data revealed that B12 supplementation, in aged female mice, significantly impacted the expression in mitochondria-related proteins in the muscle and the majority of mitochondrial-related proteins, that were impacted, had increased expression. Further, functional annotation demonstrated that mitochondrial proteins were related to the TCA cycle, OXPHOS, and oxidative stress. Collectively, these results support that B12 supplementation, in older adults, could improve skeletal muscle mitochondrial biology, a hallmark of aging.

B12 supplementation increased mitochondrial content in old muscle. It is likely that mitochondrial biogenesis is an indirect response to improvement in B12 availability, recognizing that the two enzymes to which B12 is a co-factor are both linked to mitochondrial metabolism. Because skeletal muscle is mitochondrially dense and requires a high amount of ATP for function and contraction, increased or preserved mitochondrial function has potential to greatly impact skeletal muscle physiology. We showed that CS activity, which is seen to decrease in aging muscle, contributing to the decline in skeletal muscle functional capacity (45), increased with B12 supplementation in aged muscle. Additionally, the observed enrichment of proteins involved in mtDNA biosynthesis and maintenance, including TFAM, along with an increase in mtDNA content, suggest that B12 may impact the mtDNA level, potentially leading to enhanced mitochondrial function. TFAM and mtDNA are seen to decline with advancing age (36). MtDNA copy number is seen to decline with advancing age (46) and shows a positive relationship to B12 levels in blood of older adults (46). TFAM has been observed to protect mtDNA, leading to an increase in mitochondrial function in skeletal muscle, in states of disuse (37). Increased expression of TFAM could be seen as a potential alleviator of sarcopenia by supporting mitochondrial quality control and function (37, 47). As we have seen increased TFAM with B12 supplementation, this could indicate that B12 can alleviate mtDNA damage and promote mitochondrial function, thus having potential to alleviate a potential contributor to sarcopenia.

Twelve weeks of B12 supplementation had a positive impact on mitochondrial morphology in older female skeletal muscle. Conversely, others have demonstrated that the skeletal muscle and cardiac tissue of B12 deficient young sheep (18) and the skeletal muscle of older adults (13) have distorted mitochondrial morphology. In line with impairments in B12 metabolism impacting mitochondrial morphology, humans and mice with genetic deficiency of MMUT have distorted mitochondria, including cristae with reduced matrix density and rarefication (48). It is possible that the observed improvements in morphology are due to increased MMUT activity; we observed a trend toward reduced MMA in the SUPP mice (vs. CNT), supporting the notion that MMUT activity is improved with B12 supplementation in the old female muscle. Intriguingly, we also identified an increase in MMUT associated proteins, including citramalyl-CoA lyase (CLYBL) [1.8-fold increase]. CLYBL is responsible for degrading itaconate, a MMUT inhibitor that is widely associated with decreased B12 levels (49). Further exploration is necessary to understand the direct impact of MMUT activity on mitochondrial morphological dynamics.

It is also possible that improvements in mitochondrial morphology are mediated by reductions in ROS levels. Elevated ROS levels have been associated with declines in mitochondrial morphology, including increased fragmentation and mitophagy (50). We did not observe any changes in the major protein complexes that are known to support cristae biogenesis (i.e., OPA1, F1F0 ATP Synthase, MICOS), suggesting that morphological improvements may instead arise from a more favorable redox environment rather than direct remodeling.

Increased abundance of TCA cycle proteins is critical, as they generate the reducing equivalents and metabolites required to fuel the ETC and support OXPHOS and ATP production (51). Despite increased expression of complex I and IV proteins with B12 supplementation and previously observed increased complex I and IV OCR in B12 supplemented, aged male mice (19), we did not observe an improvement in mitochondrial complex activity (i.e., oxygen consumption rate) in the aged female mice. Differences could be due to duration of supplementation: 12 weeks in the current study vs. 8 weeks (19). This could also be due to sex differences in age-related mitochondrial deterioration (52).

### Study limitations

There were a few study limitations. Physical activity was not assessed with this study yet could provide explanation for mitochondrial biogenesis and increased muscle weight. Mitochondrial biogenesis occurs with aerobic exercise training (53, 54), to meet energy and metabolic needs (55). Potential changes in physical activity and energetics are of interest in future studies. Another limitation was the human cell lines, which was from a single donor. This study is focused on the impact of B12 supplementation on aging skeletal muscle, in which the cell model is used only as a guide for potential mechanism and pathway that B12 may utilize within aging muscle. Within the cell model, mechanisms of aging were not accounted for, which could be considered a minor limitation.

## Conclusions

Our results indicate that in aged female mice, 12 weeks of B12 supplementation improved skeletal muscle mitochondrial biology, specifically improving mitochondrial DNA, content, morphology and proteins. Because mitochondria are the primary energy producers within the cell, declines in their function and efficiency can have detrimental consequences to overall tissue function (56). Despite this critical role, and the negative impact previously noted of B12 deficiency in relation to mitochondrial health, the effects of B12 on skeletal muscle remain underexplored and, to our knowledge, have not been mechanistically evaluated. These results, suggest that B12 supplementation may be a viable therapy to mitigate age-related mitochondrial degeneration within skeletal muscle.

## ACKNOWLEDGMENTS

The authors responsibilities are as follows. ARW, MSF, AET: research design; ARW, WM, LFC, AY, SRS, LEO, CTP, SR, JM, MSF, AET: performed experiments and analyzed data; ARW, AY, WM, JM, MSF, AET: drafted and edited paper; all authors reviewed the final manuscript.

