## Supplemental Figures for "Vitamin B12 Improves Skeletal Muscle Mitochondrial Biology in Aged Mice"

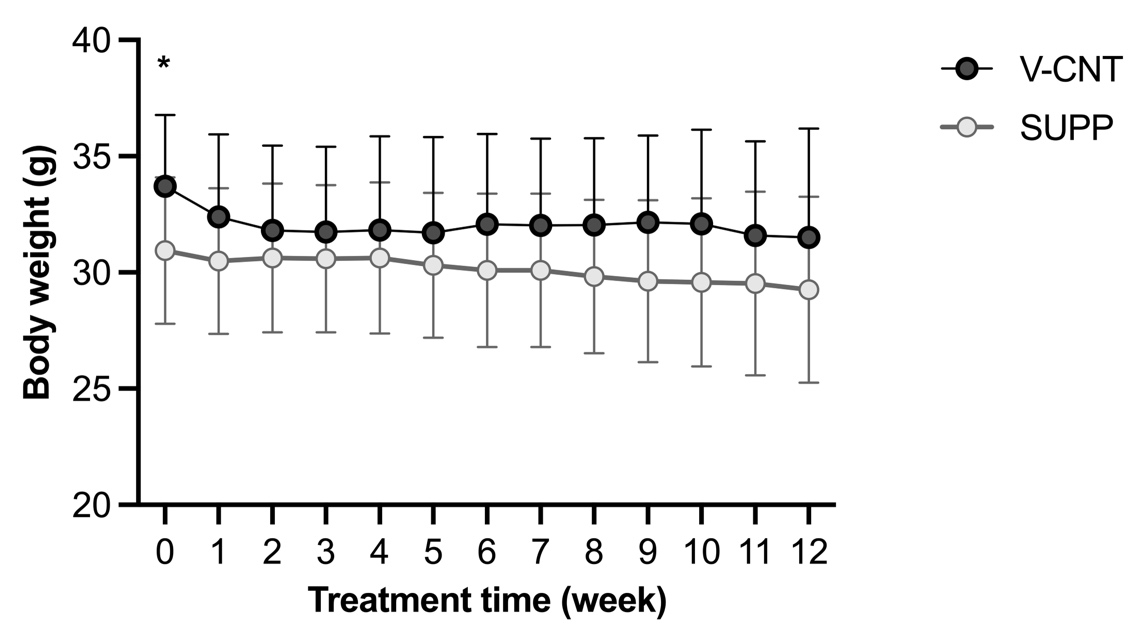

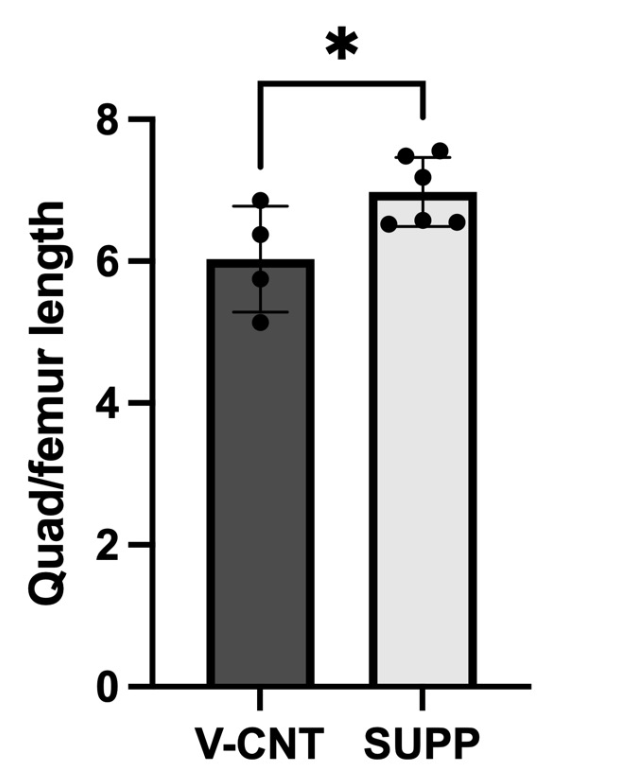

**A**

**C**

**B**

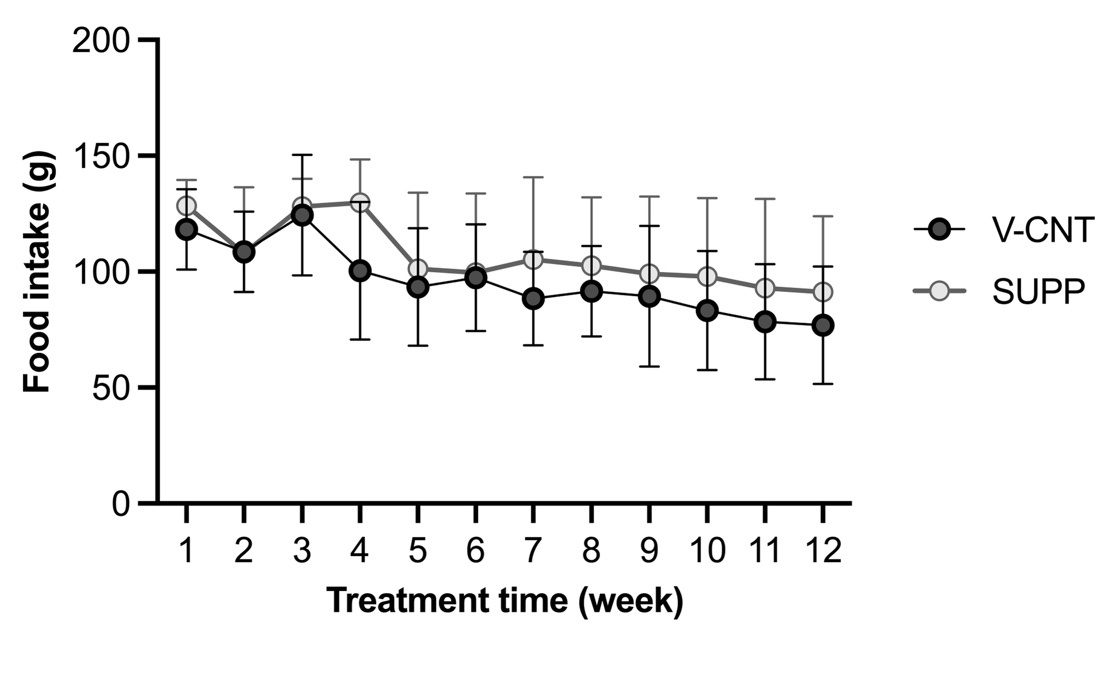

**SFig. 1. B12 treatment impacts muscle mass independent of body mass. A.** Body weight was non-significantly changed within treatment groups over the course of 12-week treatment. **B.** Food consumption was non-significantly different within treatment groups over the course of the 12-week treatment. **C.** B12 supplemented female mice displayed significantly greater muscle mass compared to vehicle treated. *p<0.05

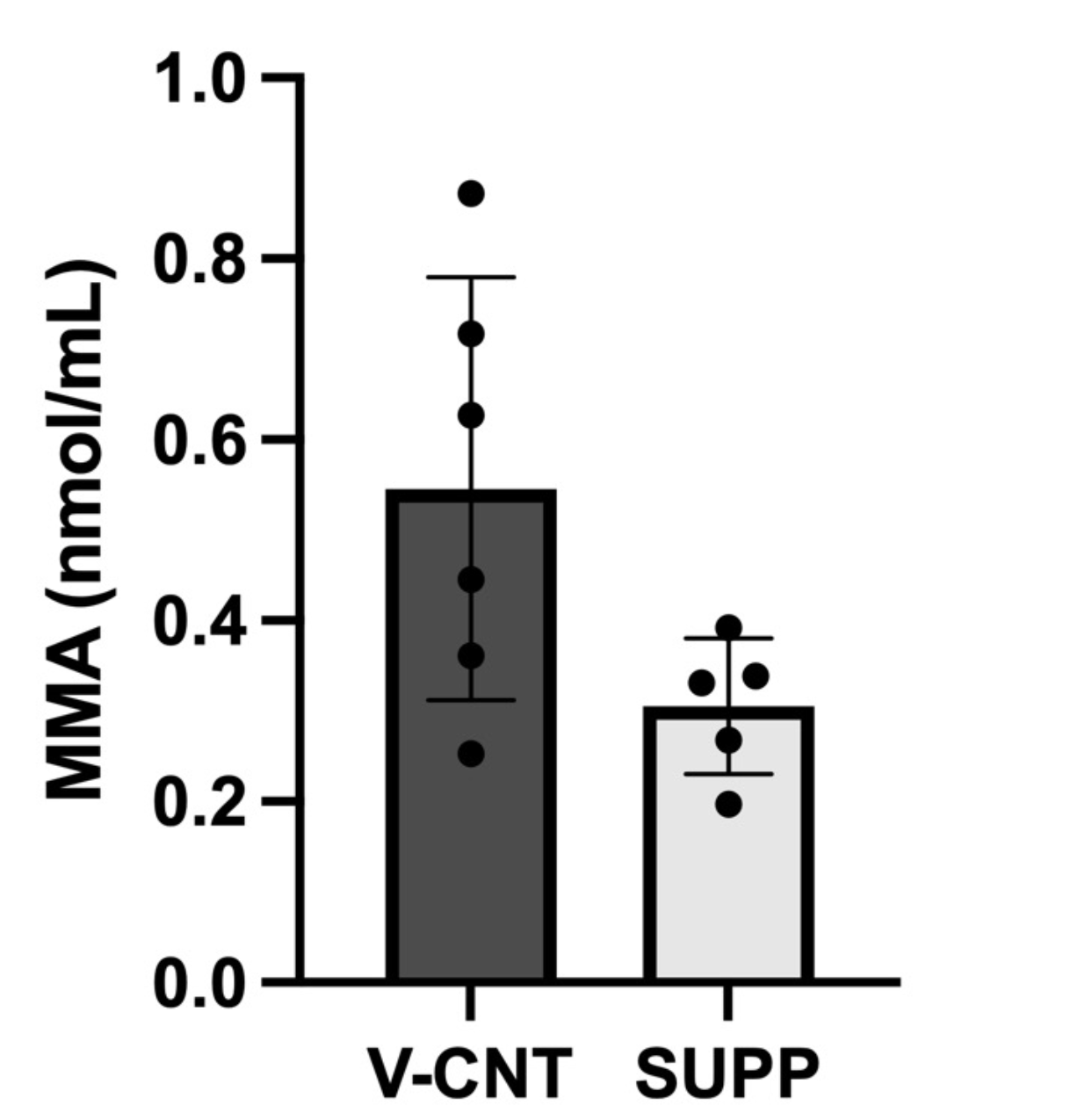

**SFig. 2. Methylmalonic acid shows trend towards decrease with IM B12 injection treatment.** MMA shows almost significant decrease (p=0.056) in B12 treated mice (n=10) compared to vehicle (n=8)

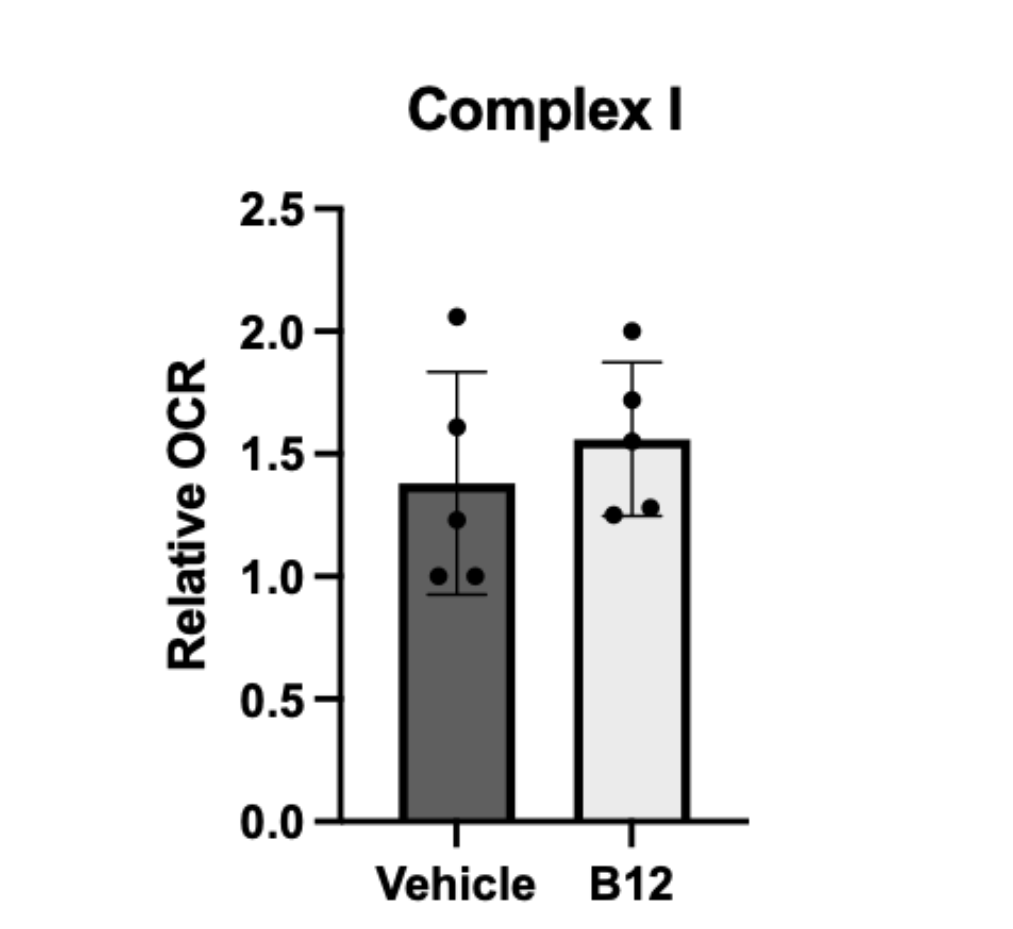

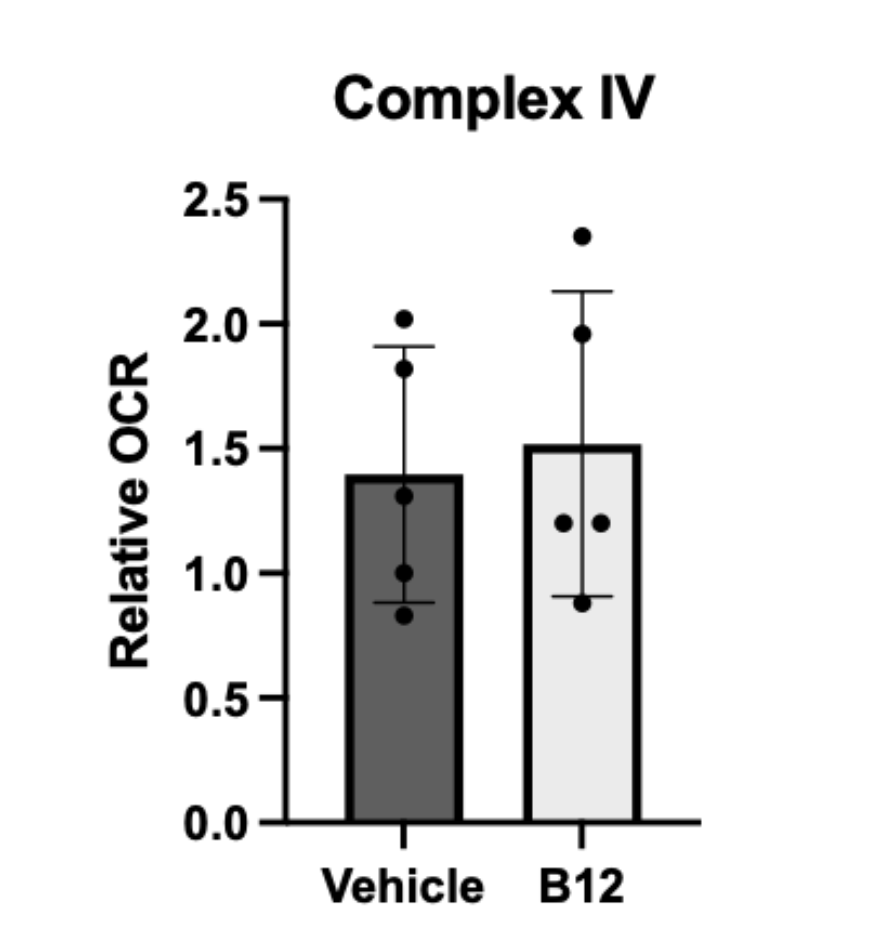

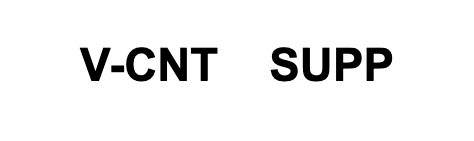

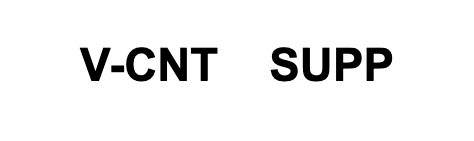

**SFig. 3. Mitochondrial oxygen consumption was unaffected by B12 supplementation in aged female mice.** No difference was detected in maximal oxygen consumption in Complex I or Complex IV in gastrocnemius of aged female mice between vehicle (n=8) and B12 supplemented (n=10) groups
